# Electrophysiological resting-state signatures link polygenic scores to general intelligence

**DOI:** 10.1101/2025.01.17.633388

**Authors:** Rebecca Engler, Christina Stammen, Stefan Arnau, Javier Schneider Penate, Dorothea Metzen, Jan Digutsch, Patrick D. Gajewski, Stephan Getzmann, Christoph Fraenz, Jörg Reinders, Fabian Streit, Sebastian Ocklenburg, Daniel Schneider, Michael Burke, Jan G. Hengstler, Carsten Watzl, Michael A. Nitsche, Robert Kumsta, Edmund Wascher, Erhan Genꞔ

## Abstract

Intelligence is associated with important life outcomes. Behavioral, genetic, structural, and functional brain correlates of intelligence have been studied for decades, but questions remain as to how genetics are related to trait expression and what intermediary role brain properties play. This study investigated these mediations in a representative sample of 434 individuals, comprising young and older adults. Polygenic scores (PGS) for intelligence were calculated. Resting-state EEG recordings were analyzed using graph theory quantifying functional connectivity across different frequencies. We tested whether global and local graph metrics like efficiency and clustering mediated the association between PGS and intelligence. PGS significantly predicted variance in intelligence and were related to frequency-specific graph metrics in areas predominantly located in parieto-frontal regions, which in turn were associated with intelligence. These findings, which are based on the first study linking PGS to intelligence using EEG-derived graph metrics, advance our understanding of the neurogenetics of intelligence.

## Introduction

Interindividual differences in human intelligence and their neural basis have fascinated researchers for over a century. Intelligence is commonly defined as „a very general mental capability that, among other things, involves the ability to reason, plan, solve problems, think abstractly, comprehend complex ideas, learn quickly, and learn from experience. It is not merely book learning, a narrow academic skill, or test-taking smarts. Rather, it reflects a broader and deeper capability for comprehending our surroundings – ‘catching on’, ‘making sense’ of things, or ‘figuring out’ what to do” ^1^. It is one of the most stable psychological traits ^2,3^ and has been found to correlate with important life outcomes like health ^4^, occupation ^5^, educational attainment ^6^, and mortality ^2^. Even though behavioral, genetic as well as structural, and functional brain correlates of human intelligence have been studied for decades, the exact neurogenetic mechanisms behind intelligent thinking are still hardly understood.

Twin studies revealed that genetic factors can explain about 50% of interindividual differences in intelligence, making intelligence a highly heritable trait ^7,8^. On the molecular level, genome-wide association studies (GWAS) investigated the association of single-nucleotide polymorphisms (SNP) – which are changes in a single base pair on the genome – with intelligence ^9–11^. They revealed hundreds of SNPs significantly associated with intelligence and identified intelligence as a highly polygenic trait. Using the summary statistics of GWAS, it is possible to calculate a polygenic score (PGS), which summarizes the predisposition of a participant for a certain phenotype ^12^. PGS for intelligence are able to predict up to 7% of intelligence in independent validation samples ^7,13^.

The most likely pathway through which SNPs influence intelligent thinking involves structural and functional brain properties. A prominent neuroscientific model of intelligence is the Parieto-Frontal Integration Theory (P-FIT). Originally postulated by Jung and Haier (2007), this model states that intelligent thinking relies on efficient information transfer between brain regions across the cortex, with a crucial role for frontal and parietal cortices. Interestingly, while the P-FIT underlines the importance of efficient information transfer, the original studies did not include research concerning brain connectivity. One method of quantifying connectivity in neuroscience employs graph theory. A graph consists of nodes, which are spatially defined in the brain, and edges, which represent connections between these nodes ^15^. Two well-researched graph-theoretical metrics are efficiency, which quantifies how efficiently information is transferred between brain areas, and clustering, which quantifies the cliquishness of a brain network. Studies employing diffusion-weighted imaging (DWI) have identified that the brain’s global efficiency is positively related to intelligence ^16–21^. While some of these studies report associations to be strongest with nodes in the P-FIT network ^17,19^, other studies identified primarily areas outside of the P-FIT network ^18^. Also the clustering coefficient has been reported to be positively associated with intelligence ^20^, but other studies did not find this association ^19^.

Resting-state functional magnetic resonance imaging (rsfMRI) studies investigating the relation between graph metrics and intelligence have come to inconclusive results. Van den Heuvel et al.^22^ were the first to report a positive association between intelligence and global efficiency, however, this effect could not be replicated in subsequent studies ^23–25^. On a regional level, Pamplona et al. ^25^ mainly reported areas associated with intelligence outside of P-FIT, while Hilger et al. ^23^ identified some P-FIT regions. A recent multicenter study ^26^ found no robust associations between intelligence and global or regional graph metrics, concluding that static rsfMRI in combination with graph theory is not suited for investigating the neural underpinnings of intelligence.

One often-overlooked alternative to investigate the functional connectome is electroencephalography (EEG). Network Neuroscience Theory argues that intelligence is not only sustained by efficiency but also by the flexibility of brain networks ^27^. EEG, with its superior temporal resolution, might be better suited for understanding the neural mechanisms behind intelligence despite its lower spatial resolution compared to fMRI. EEG data can be segmented into different temporal frequencies, allowing for performing highly differentiated connectivity analyses for specific frequencies.

Very few studies have investigated intelligence and its relation to resting-state EEG (rsEEG), exclusively conducted in young adults. Langer et al. ^28^ found that in male adults the EEG alpha band clustering and efficiency was positively associated with intelligence. Additionally, the parietal cortex was identified as a main hub of the rsEEG network. Subsequent studies, however, found contradictory results on the association between efficiency and clustering in EEG frequency bands and intelligence ^29,30^. The three studies differ in mean age and sex distribution, affecting result comparability. Large-scale cohort studies using rsEEG graph metrics are needed to better understand the association between the functional connectome and intelligence.

Most rsfMRI and rsEEG studies have focused on children ^22^, young adults ^23,25,26^, or mixed samples ^26^, with no study investigating functional graph metrics related to intelligence exclusively in older adults. Since the human connectome changes with age ^31^ investigating older adults bridges a gap in deciphering the neural underpinnings of intelligence.

Another often overlooked factor is the genetic basis of the functional connectome. Interestingly, the heritability of rsEEG connectivity metrics has been analyzed in a twin study. Smit et al. ^32^ reported that across different frequency bands, 46-89% of individual differences in the clustering coefficient are heritable, suggesting that functional connectivity patterns have a stable biological basis and might be biomarkers of individual differences in brain functioning. Several other studies also support the heritability of different EEG measures ^33–38^.

While the above-mentioned studies investigated the heritability and molecular genetic basis of EEG derived functional brain organization, the current study goes a step further, investigating the triad of molecular genetics, intrinsic electrophysiological connectivity and human intelligence.

Several studies on structural brain properties ^39–41^ found that the surface area of parietal and frontal areas and the structural connectivity of frontal areas mediate between genetic variation and intelligence. Only one study has investigated the mediating effect of rsfMRI functional graph metrics ^40^, finding no brain areas where nodal efficiency mediated the effect of SNPs on intelligence, likely because rsfMRI nodal efficiency is not robustly associated with intelligence ^26^. As mentioned before, rsEEG may offer some advantages over rsfMRI, making it potentially better suited for investigating how genetics connect to intelligence. With this regard, exploratory mediation analyses were employed in this study to investigate whether frequency-specific connectivity at a global level or at a regional level acts as a mediator between PGS of general intelligence (PGS_GI_) and intelligence. To investigate the robustness of the rsEEG connectivity properties, test-retest reliability of global and regional graph metrics was determined by means of intra-class correlation (ICC). An illustration of the pre-processing and analysis strategy is depicted in Figure 1.

**Figure 1.**
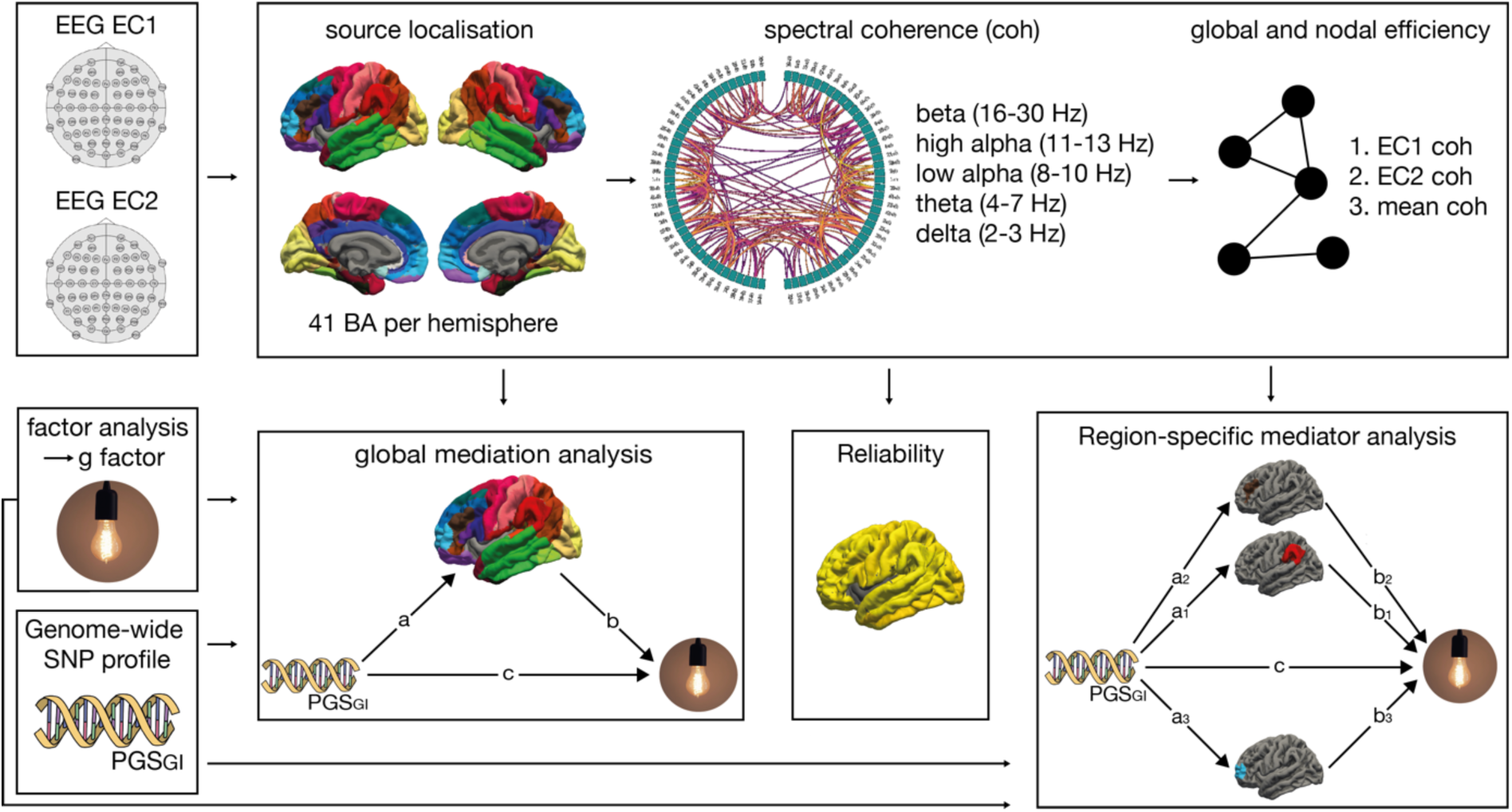
Pre-processing and analysis strategy of the data sets. First, source localization was performed for the two electroencephalography (EEG) resting-state recordings with eyes closed (EC1 and EC2). Per hemisphere, 41 regions of interest (ROI) were defined, corresponding to 41 Brodmann areas (BA). Second, all-to-all inter-ROI functional connectivity was calculated as the spectral coherence (coh) for the beta (16-30 Hz), high alpha (11-13 Hz), low alpha (8-10 Hz), theta (4-7 Hz), and delta (1-3 Hz) frequency range for EC1 and EC2, resulting in two 82 x 82 resting-state networks per frequency band. Third, the global and nodal efficiency of EC1 and EC2 were calculated. EC1 coh and EC2 coh were used to calculate mean coherence (mean coh). Fourth, a factor analysis was carried out to extract *g*, the factor of general intelligence. Fifth, the polygenic score for general intelligence (PGS_GI_) was calculated. Sixth, graph theoretical metrics on the global (global efficiency, global clustering) and nodal level (nodal efficiency, local clustering) were computed and global as well as region-specific mediation analyses were performed with PGS_GI_ as the independent variable, *g* as the dependent variable, and the graph theoretical metrics as respective mediators. Finally, the reliability of the graph’s theoretical metrics was defined by calculating the intra-class correlation (ICC) of the global efficiency and global clustering and of the nodal efficiency and local clustering of each frequency band.

The present mediation study consisting of data of 434 individuals is, to the best of our knowledge, the largest dataset to investigate rsEEG graph metrics and their relation to intelligence. To account for age-related connectome changes, the sample was divided into young (n = 199, aged 20 – 40 years) and older adults (n = 235, aged 40 – 70 years). Ultimately, this study is the first mediation study to investigate whether rsEEG graph metrics are the missing link to how genes shape intelligence.

## Results

### Partial correlations

Partial correlations between *g*, global efficiency, and global clustering coefficient and partial correlations between the PGS_GI,_ global efficiency, and global clustering for young and older adults are shown in the Supplementary Table S1 and S1. No correlation reached statistical significance (.097 ≤ *p* ≤ .999). Results for the whole sample can also be found in Supplementary Tables S1 and S2.

### Global mediation analysis

Results of the global mediation analyses are shown in Supplementary Table S3 for young adults, older adults, and the full sample. None of the global mediation analyses reached statistical significance. Even though partial correlations conducted prior to this analysis did not reveal a significant effect, the global mediation analysis was still conducted, as the statistical significance of paths a and b alone is not a precondition for a significant mediation effect ^42,43^.

### Brain area-specific mediation analyses

#### Nodal efficiency in young adults

Results of the brain area-specific mediation analysis for nodal efficiency in young adults are depicted in Figure 2. All effect sizes with the respective Brodmann areas are available in Supplementary Table S4, S5, and S6.

**Figure 2.**
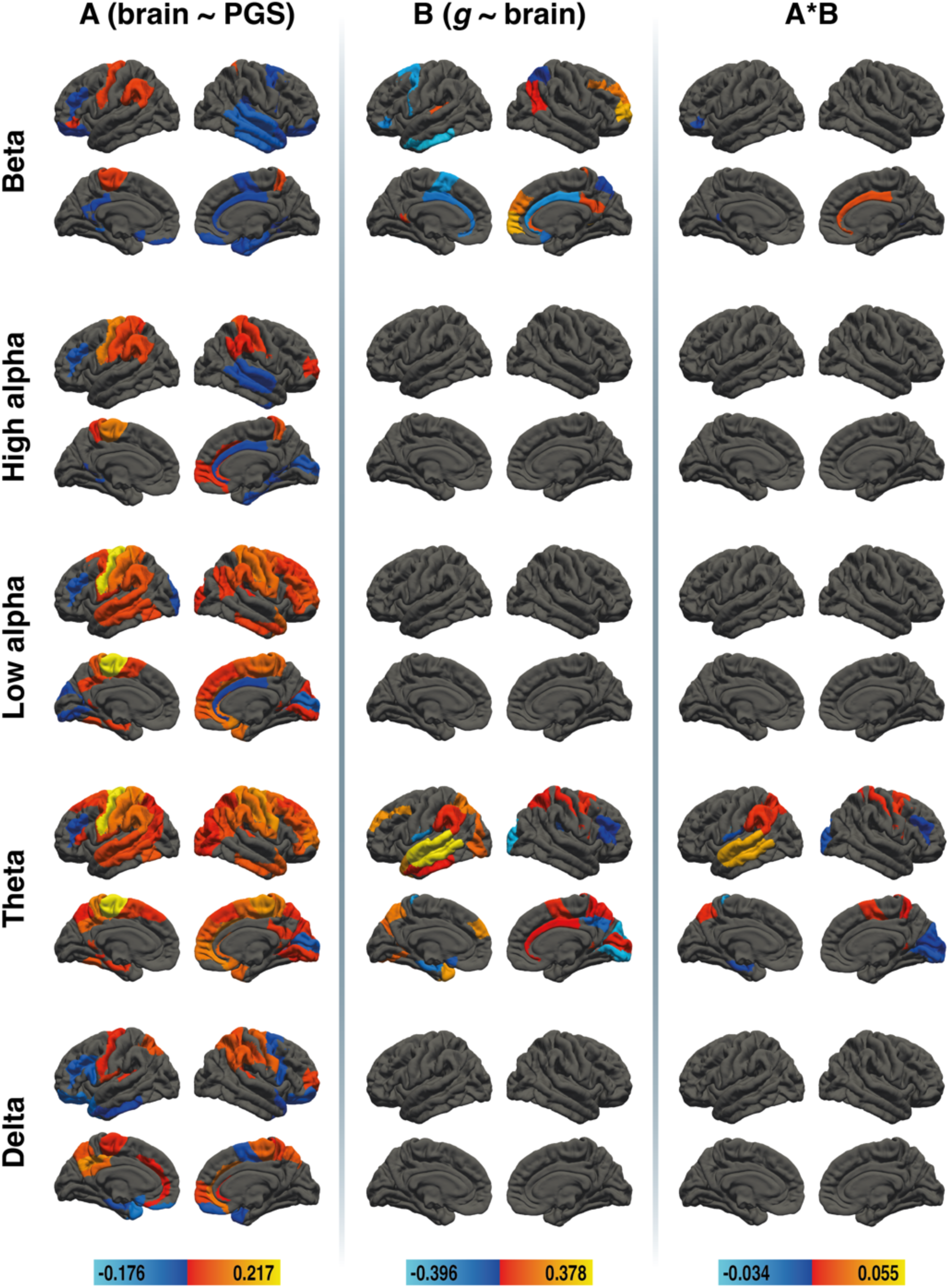
Results of the region-specific mediation analysis via elastic-net regression (nodal efficiency) in young adults. Respective mediators were 41 cortical areas in the left and 41 cortical areas in the right hemisphere of five different frequency bands (delta, theta, low alpha, high alpha, and beta, from bottom to top). The figure shows the results from path a analysis (brain ∼ PGS), path b analysis (*g* ∼ brain), and the mediation effect (from left to right). Brain surfaces are shown in lateral and sagittal views, for the left and right hemispheres. Positive effects are depicted in red and yellow, negative effects are depicted in blue. A full list of all effect sizes is available in Supplementary Table S4, Supplementary Table S5, and Supplementary Table S6.

#### Beta

PGS_GI_ was associated with beta-frequency nodal efficiency in 25 cortical areas, 5 showed a positive association, and 20 showed a negative association. Beta-frequency nodal efficiency was associated with *g* in 17 areas, 10 showed a positive association, and 7 showed a negative association. The beta-frequency nodal efficiency of right BA24 showed a positive mediation of the effect of PGS_GI_ on *g*, and left BA26, left BA47 and right BA30 showed a negative mediation effect.

#### High Alpha

PGS_GI_ was associated with high alpha-frequency nodal efficiency in 26 cortical areas, while no areas were associated with *g*, resulting in no selected mediators.

#### Low alpha

PGS_GI_ was associated with low alpha-frequency nodal efficiency in 42 cortical areas, while no areas were associated with *g*, *resulting in* no selected mediators.

#### Theta

PGS_GI_ was associated with theta-frequency nodal efficiency in 47 cortical areas, 45 showed a positive association, and 2 showed a negative association. Theta-frequency nodal efficiency was associated with *g* in 24 areas, 14 showed a positive association, and 10 showed a negative association. The theta-frequency nodal efficiency of left BA7, BA22, BA40 as well as right BA1, BA5, BA6, and BA26 showed a positive mediation of the effect of PGS_GI_ on *g*, and left BA3, BA33, BA36, and BA43 as well as right BA17, BA18, and BA46 showed a negative mediation effect.

#### Delta

PGS_GI_ was associated with delta-frequency nodal efficiency in 29 cortical areas, while no areas were associated with *g*, resulting in no selected mediators.

#### Nodal efficiency in older adults

Results for the brain-area-specific mediation analysis for nodal efficiency in older adults are depicted in Figure 3. All effect sizes with the respective Brodmann areas are available in Supplementary Table S7, S8, and S9.

**Figure 3.**
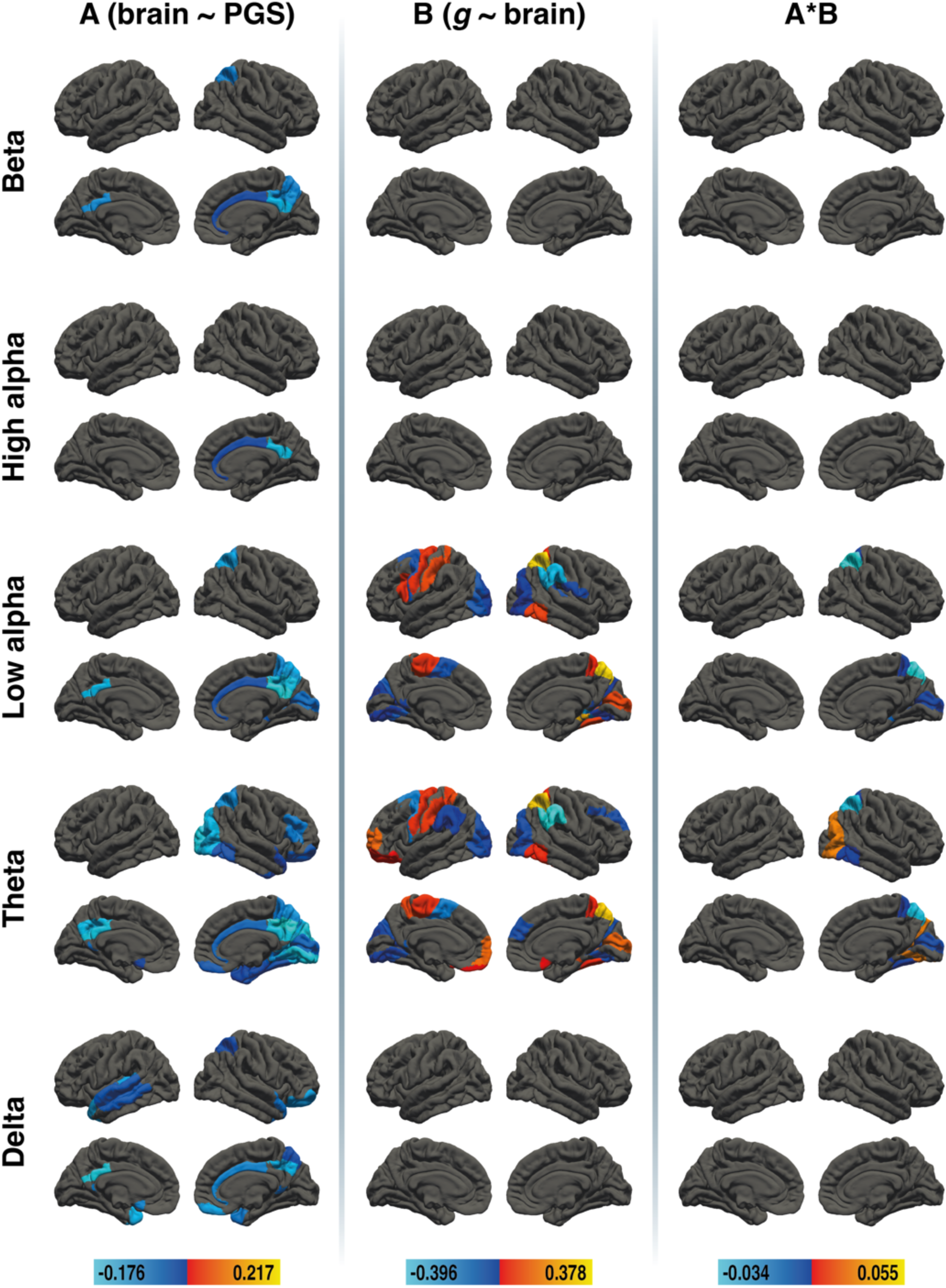
Results of the region-specific mediation analysis via elastic-net regression (nodal efficiency) in older adults. Respective mediators were 41 cortical areas in the left and 41 cortical areas in the right hemisphere of five different frequency bands (delta, theta, low alpha, high alpha, and beta, from bottom to top). The figure shows the results from path a analysis (brain ∼ PGS), path b analysis (*g* ∼ brain), and the mediation effect (from left to right). Brain surfaces are shown in lateral and sagittal views, for the left and right hemispheres. Positive effects are depicted in red and yellow, negative effects are depicted in blue. A full list of all effect sizes is available in Supplementary Table S7, Supplementary Table S8, and Supplementary Table S9.

#### Beta

PGS_GI_ was associated with beta-frequency nodal efficiency in 5 cortical areas, while no areas were associated with *g*, resulting in no selected mediators.

#### High alpha

PGS_GI_ was associated with high alpha-frequency nodal efficiency in 2 cortical areas, while no areas were associated with *g*, resulting in no selected mediators.

#### Low alpha

PGS_GI_ was associated with low alpha-frequency nodal efficiency in 8 cortical areas, all showed a negative association. Low alpha-frequency nodal efficiency was associated with *g* in 21 areas, 10 showed a positive association, and 11 showed a negative association. The low alpha-frequency nodal efficiency of right BA5, BA7, BA17, and BA28 showed a negative mediation of the effect of PGS_GI_ on *g*, and no area showed a positive mediation effect.

#### Theta

PGS_GI_ was associated with theta-frequency nodal efficiency in 24 cortical areas, all showed a negative association. Theta-frequency nodal efficiency was associated with *g* in 20 areas, 12 showed a positive association, and 8 showed a negative association.

The theta-frequency nodal efficiency of right BA19 showed a positive mediation of the effect of PGS_GI_ on *g*, and right BA5, BA7, BA17, BA28, and BA37 showed a negative mediation effect.

#### Delta

PGS_GI_ was associated with delta-frequency nodal efficiency in 13 cortical areas, while no areas were associated with *g*, resulting in no selected mediators.

#### Local clustering in young adults

Results for the brain-area-specific mediation analysis for local clustering in young adults are depicted in Figure 4. All effect sizes with the respective Brodmann areas are available in Supplementary Table S10, S11, and S12.

**Figure 4.**
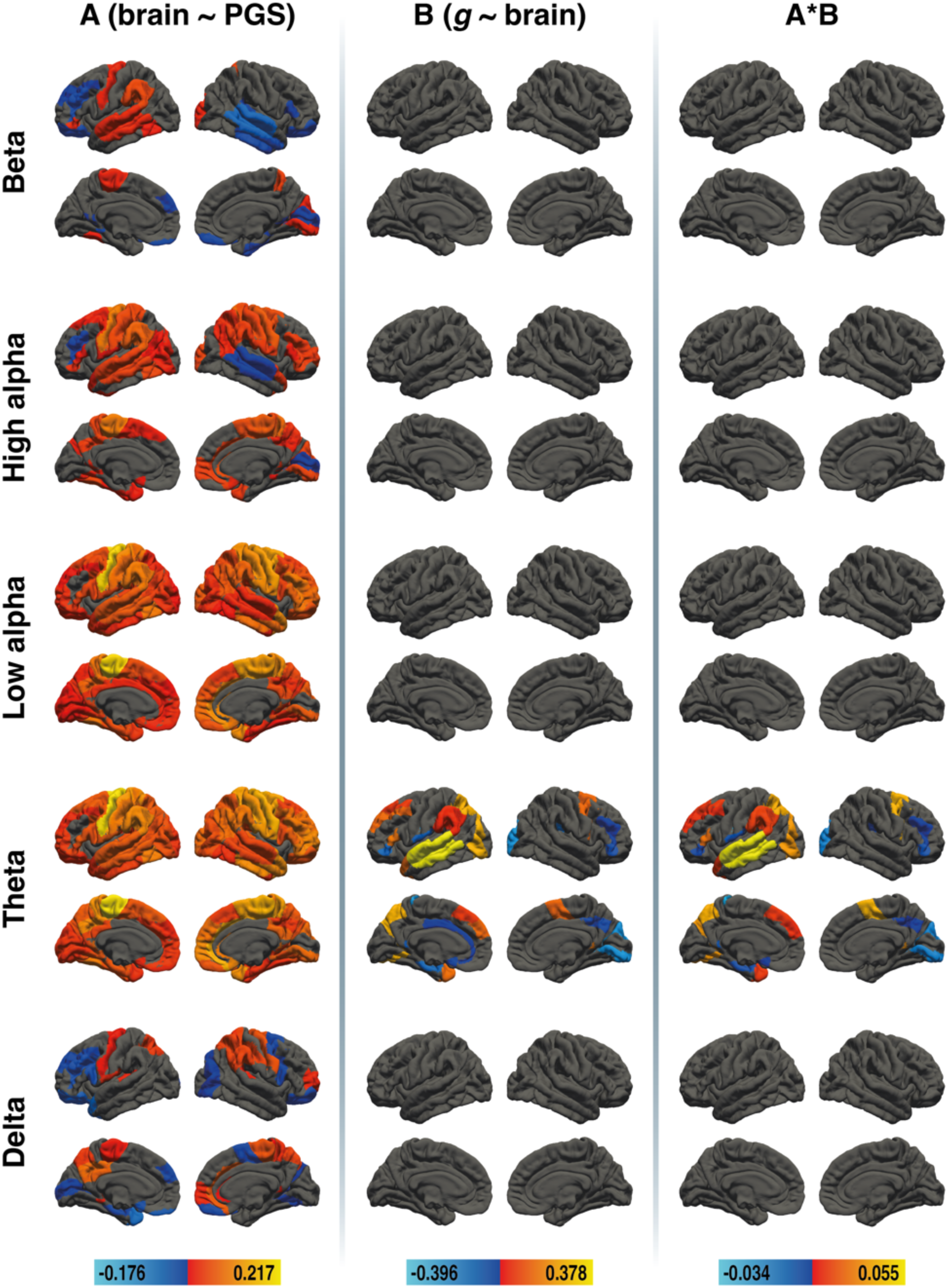
Results of the region-specific mediation analysis via elastic-net regression (local clustering) in young adults. Respective mediators were 41 cortical areas in the left and 41 cortical areas in the right hemisphere of five different frequency bands (delta, theta, low alpha, high alpha, and beta, from bottom to top). The figure shows the results from path a analysis (brain ∼ PGS), path b analysis (*g* ∼ brain), and the mediation effect (from left to right). Brain surfaces are shown in lateral and sagittal views, for the left and right hemispheres. Positive effects are depicted in red and yellow, negative effects are depicted in blue. A full list of all effect sizes is available in Supplementary Table S10, Supplementary Table S11, and Supplementary Table S12.

#### Beta

PGS_GI_ was associated with beta-frequency local clustering in 24 cortical areas, while no areas were associated with *g*, resulting in no selected mediators.

#### High alpha

PGS_GI_ was associated with high alpha-frequency local clustering in 48 cortical areas, while no areas were associated with *g*, resulting in no selected mediators.

#### Low alpha

PGS_GI_ was associated with low alpha-frequency local clustering in 78 cortical areas, all showed a positive association. There were no areas whose low alpha-frequency local clustering was associated with *g* and thus no selected mediators.

#### Theta

PGS_GI_ was associated with theta-frequency local clustering in 78 cortical areas, all showed a positive association. Theta-frequency nodal efficiency was associated with *g* in 23 areas, 10 showed a positive association, and 13 showed a negative association. The theta-frequency local clustering of left BA7, BA8, BA9, BA19, BA22, BA38, BA40 and BA45 as well as right BA6 and BA26 showed a positive mediation of the effect of PGS_GI_ on *g*, and left BA3, BA25, BA26, BA33, BA36, BA43, and BA47 as well as right BA18, BA31, BA41, BA46 and BA47 showed a negative mediation effect.

#### Delta

PGS_GI_ was associated with delta-frequency local clustering in 31 cortical areas, while no areas were associated with *g*, resulting in no selected mediators.

#### Local clustering in older adults

Results for the brain-area-specific mediation analysis for local clustering in older adults are depicted in Figure 5. All effect sizes with the respective Brodmann areas are available in Supplementary Table S13, and S14.

**Figure 5.**
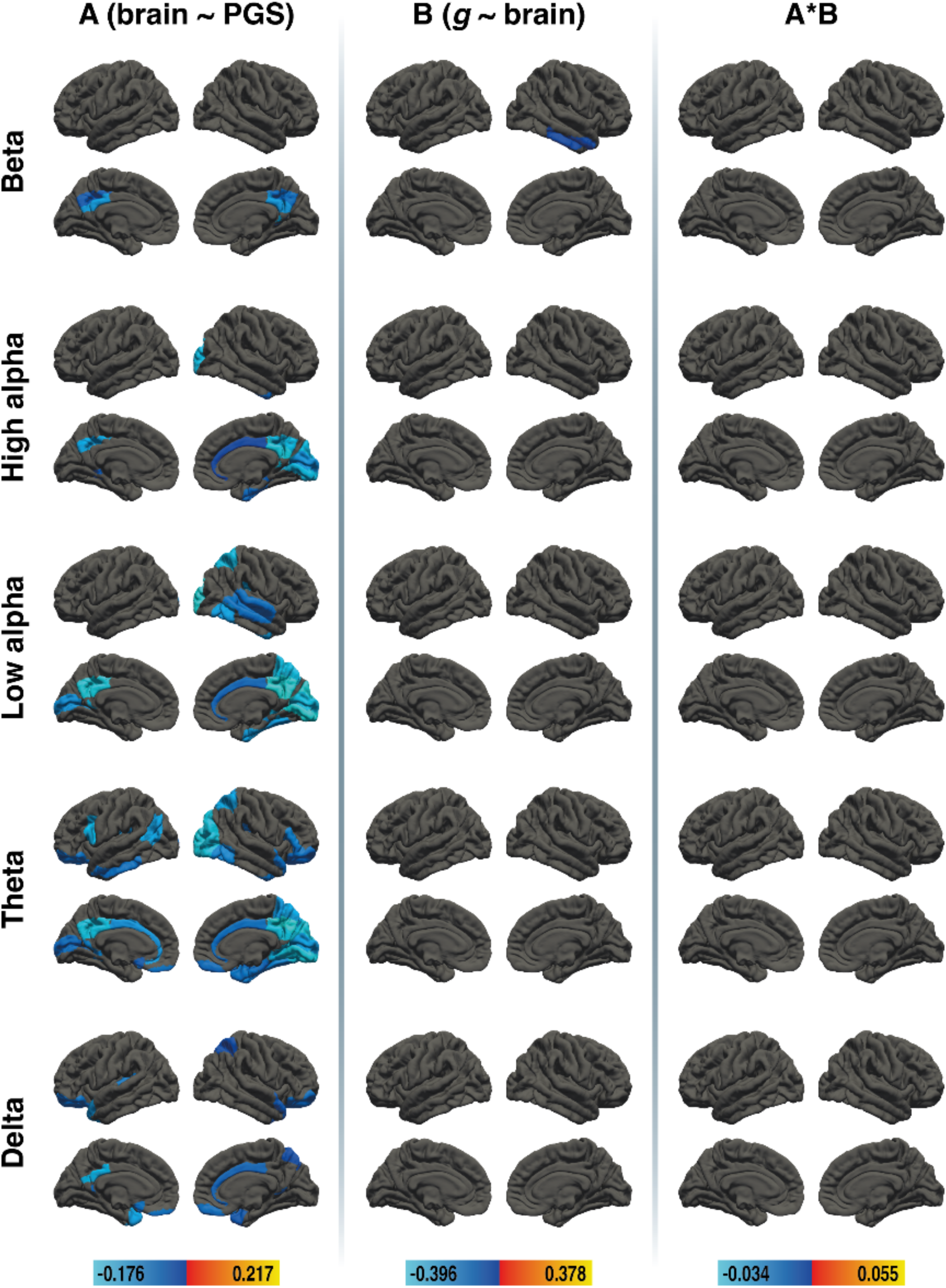
Results of the region-specific mediation analysis via elastic-net regression (local clustering) in older adults. Respective mediators were 41 cortical areas in the left and 41 cortical areas in the right hemisphere of five different frequency bands (delta, theta, low alpha, high alpha, and beta, from bottom to top). The figure shows the results from path a analysis (brain ∼ PGS), path b analysis (*g* ∼ brain), and the mediation effect (from left to right). Brain surfaces are shown in lateral and sagittal views, for the left and right hemispheres. Positive effects are depicted in red and yellow, negative effects are depicted in blue. A full list of all effect sizes is available in Supplementary Table S13 and Supplementary Table S14.

#### Beta

PGS_GI_ was associated with beta-frequency nodal efficiency in 6 cortical areas, while one area was associated with *g*, but not with PGS_GI_, resulting in no selected mediators.

#### High alpha

PGS_GI_ was associated with high alpha-frequency nodal efficiency in 10 cortical areas, while no areas were associated with *g*, resulting in no selected mediators.

#### Low alpha

PGS_GI_ was associated with low alpha-frequency nodal efficiency in 17 cortical areas, while no areas were associated with *g*, resulting in no selected mediators.

#### Theta

PGS_GI_ was associated with theta-frequency nodal efficiency in 34 cortical areas, while no areas were associated with *g*, resulting in no selected mediators.

#### Delta

PGS_GI_ was associated with delta-frequency nodal efficiency in 11 cortical areas, while no areas were associated with *g*, resulting in no selected mediators.

### Reliability

Table 3 shows the ICC of global efficiency and global clustering coefficient for all frequency bands in young and older adults. In both groups, the reliability of global efficiency in the delta, theta, and low alpha bands can be rated as good, and global efficiency of the high alpha and beta bands can be described as excellent ^44^. The reliability of the global clustering coefficient can be rated as good for all frequency bands in both groups.

**Table 3.**
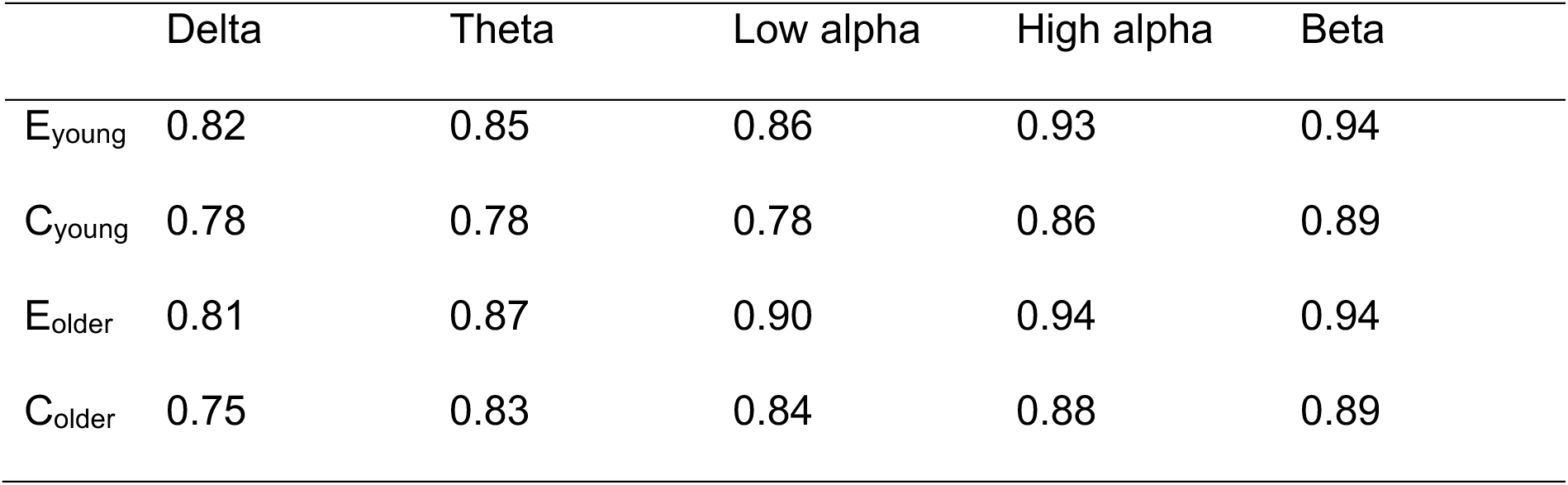
Test-retest reliability of global graph metrics for global efficiency (E) and the global clustering coefficient (C) as measured by ICC in young (E_young_, C_young_) and older adults (E_older_, C_older_). All effects were significant at the p < .001 level.

|  | Delta | Theta | Low alpha | High alpha | Beta |
| --- | --- | --- | --- | --- | --- |
| $E_{\text{young}}$ | 0.82 | 0.85 | 0.86 | 0.93 | 0.94 |
| $C_{\text{young}}$ | 0.78 | 0.78 | 0.78 | 0.86 | 0.89 |
| $E_{\text{older}}$ | 0.81 | 0.87 | 0.90 | 0.94 | 0.94 |
| $C_{\text{older}}$ | 0.75 | 0.83 | 0.84 | 0.88 | 0.89 |

Figures 6 and 7 show the ICC of nodal efficiency and local clustering for 41 cortical areas in each hemisphere for young and older adults, respectively. A complete list of areas and ICC effect sizes can be found in Supplementary Table S15 for young adults and in Supplementary Table S16 for older adults. Results for the whole sample are in Supplementary Table S17 for global metrics, and for nodal metrics in Supplementary Table S18 and Supplementary Figure 5. The delta band was the only frequency where areas showed poor reliability (ICC < 0.5). However, this was only the case for 3.7% of areas considering local clustering, and for 9.8% or 8.5% of areas considering nodal efficiency, for young and older adults respectively. Areas with poor reliability are primarily located around the posterior cingulate cortex and precuneus.

**Figure 6.**
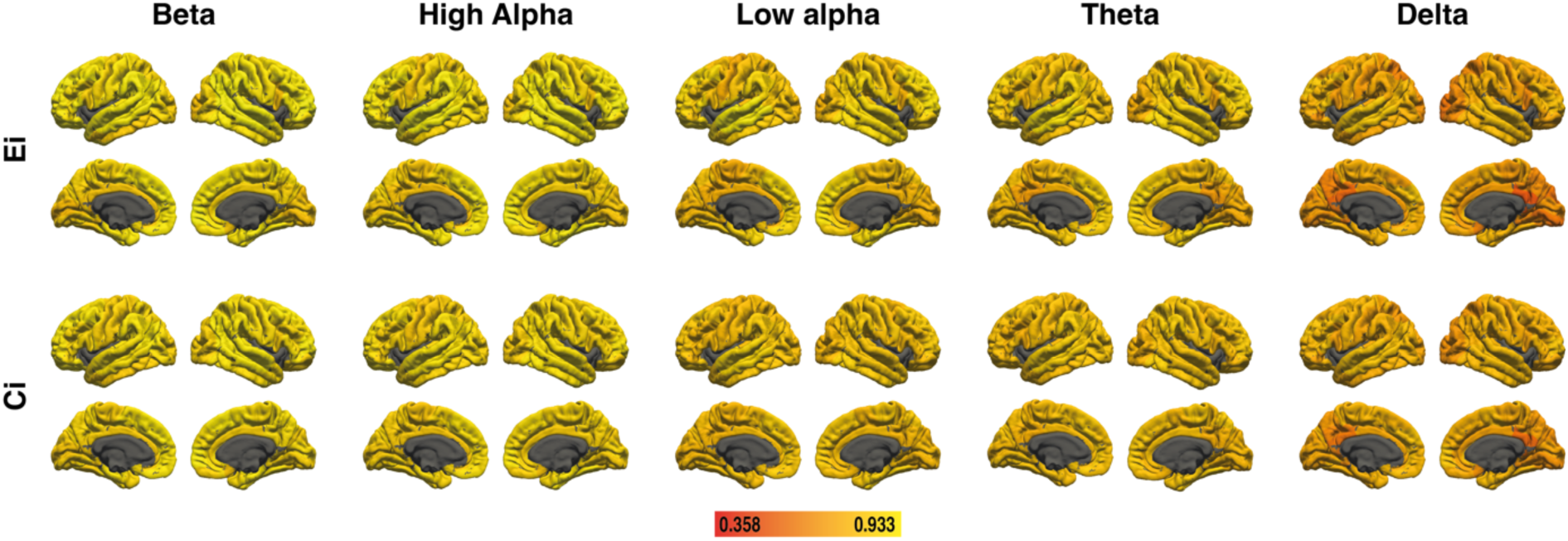
Test-retest reliability of nodal efficiency (E_i_) and local clustering (C_i_) as measured by ICC in young adults. Lighter colors indicate higher reliability.

**Figure 7.**
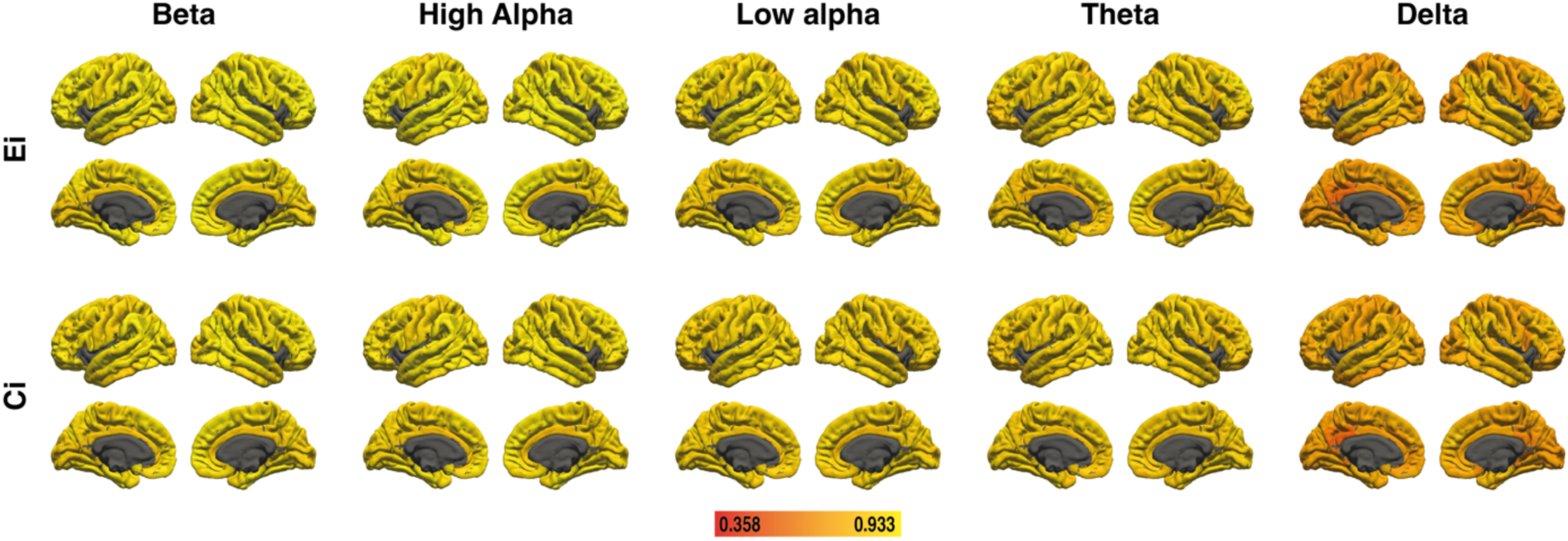
Test-retest reliability of nodal efficiency (E_i_) and local clustering (C_i_) as measured by ICC in older adults. Lighter colors indicate higher reliability.

### Results for the whole sample

All results for the whole sample can be found in the Supplementary Material (Supplementary Results section). Partial correlation coefficients between *g*, global efficiency, and the global clustering coefficient are shown in Supplementary Table S1 and between PGS_GI_, global efficiency, and the global clustering coefficient in Supplementary Table S2. Supplementary Table S3 displays the results of the global mediation analysis. Results for the brain-area-specific mediation analysis for the full sample for nodal efficiency are depicted in Supplementary Figure S3 and for local clustering in Supplementary Figure S4. Test-retest reliability of global graph metrics is shown in Supplementary Table S17. Brain-area-specific reliability results can be found in Supplementary Figure S5 and Supplementary Table S18.

## Discussion

The current study aimed to investigate how genetic variation influences intelligence by affecting the brain connectome. Using resting-state EEG data, we performed an exploratory mediation analysis to investigate the relationship between genes, graph theoretical measures, and intelligence in young and older adults. EEG, unlike fMRI, allows for frequency-specific analyses of neural activation with high temporal resolution. The study revealed brain areas with relevant frequency-specific nodal efficiency and local clustering properties in young adults, and brain areas with relevant frequency-specific nodal efficiency properties in older adults. The detected mediators carry information about the relevant brain regions and the relevant frequency bands involved in intelligent outcomes. In line with other neuroimaging studies ^23,25,26,32^, the association between PGS and intelligence was not mediated by global graph metrics. Our findings suggest that local graph metrics are suited for studying the genetic pathways influencing intelligence. Three frequency bands are of particular interest, namely beta, theta, and low alpha. With its superior temporal resolution and frequency-specificity, EEG is a promising tool for understanding the functional connectome underlying intelligence. rsEEG offers more robust connectivity measures, considering the higher reliability if compared to the same metrics for rsfMRI ^26^. The mean test-retest reliability across all frequency bands and age groups was 0.89 for global efficiency and 0.83 for global clustering. In comparison, Metzen et al.^26^ reported mean intra-class correlation coefficients for fMRI across three data sets of 0.54 for global efficiency and 0.34 for global clustering.

In young adults, beta nodal efficiency in frontal and parietal regions mediated between PGS_GI_ and intelligence. A high PGS links to increased efficiency in regions involved in semantic processing ^45,46^ and working memory ^47,48^ (ventrolateral prefrontal cortex (vlPFC)), and to decreased efficiency in regions involved in broader cognitive functions (dorsal anterior cingulate cortex (dACC) and retrosplenial cortex (RSC)) ^49,50^. Apparently, subjects with a high PGS_GI_ exhibit less efficient connectivity in regions not directly related to intelligence. The relationship between beta oscillation and intelligence is not well understood and studies on intelligence-related functions report inconsistent findings ^51^, emphasizing the need for further research.

In young adults, theta nodal efficiency in frontal, parietal, occipital, and temporal regions mediates between PGS and intelligence. A higher PGS links to higher efficiency in all regions except the primary visual cortex (V1), a low-level visual structure, which is not involved in higher-order cognitive processes. Suppression of redundant, low-level visual information is more pronounced in more intelligent individuals ^52^, which is in line with the assumption that V1 does not require to be highly efficiently connected to many other nodes within an intelligence-related network. Frontal and parietal areas are closely linked to fluid intelligence and a transcranial electrical stimulation (tES) study found higher fluid intelligence scores when theta transcranial alternating current stimulation (tACS) was applied to these regions ^53^. The effect was stronger for parietal theta tACS, which is in line with the relatively high number of parietal mediators identified in our study. Theta is involved in long-range neural communication ^54^ which might explain the wide distribution of theta mediators over the cortex. Moreover, theta plays a key role in executive control and working memory maintenance ^55–57^. For task evoked activity, it has been shown that mid-frontal theta connectivity during later stages of higher order processing in a cognitive control task explains 65% of variance in intelligence differences ^58^. Frontal and parietal regions mediate 70% of the association between genetic variants and intelligence, with a large overlap between theta-nodal efficiency and theta-local clustering regions. Theta local clustering mediated between PGS_GI_ and intelligence in areas essential for intelligence and related functions, such as the prefrontal cortex (dlPFC, mPFC) ^59,60^, Broca’s and Wernicke’s areas, along with various motor, sensory, and emotion- and memory- related regions. Taken together, theta appears to be a significant pathway for the genetic influence on intelligence.

In older adults, low alpha nodal efficiency mediated the genetic effect on intelligence in the superior parietal lobule (SPL), which is involved sensory information integration ^61^, V1, and the entorhinal cortex (EC), which is important for learning and memory ^62^.

Interestingly, no contribution of frontal areas was found. A higher PGS links to lower efficiency in V1 and EC. The fact that low alpha acts as a gateway in older adults, whereas beta mediates in young adults, might reflect an age-related shift from fast, localized communication towards a slower and broader communication. This assumption aligns with the observed age-related neural dedifferentiation, which describes a reduction of functional specificity of neural processing with aging ^63^. The aging brain decreases in within-network connectivity and increases in between-network connectivity ^64^, especially in the frontal-parietal control system and the cingulo-opercular control system, two networks that are closely linked to fluid intelligence ^27^. Theta nodal efficiency mediates the effect of PGS on general intelligence in six parietal, occipital, temporal, and occipital-temporal regions. Interestingly, four of these six regions also serve as mediators within the low alpha frequency range, suggesting that with age, fewer mediators influence intelligence but across a broader frequency range. PGS is associated with decreased theta efficiency in parts of the SPL, while in young adults the SPL is positively associated with PGS. This reversal possibly reflects age-related functional network changes. V1 consistently shows reduced nodal efficiency with high PGS in both age groups, underlining its limited role in intelligence networks.

Across the two age-groups, 44% of areas that link PGS to intelligence belong to the P-FIT model. However, 56% of all mediators are outside P-FIT, consistent with studies identifying intelligence-related brain regions beyond P-FIT ^18,40^. It is worth noting that even though morphologically certain brain areas are not associated with P-FIT, from a connectivity perspective, they still might play a role in intelligence, suggesting that intelligence is not limited to the parieto-frontal network but that a much broader neural architecture underlies human intelligence.

In total, 25 mediators were identified in young adults, and only 6 mediators were found in older adults, likely due to an age-related decrease in nodal efficiency ^31^. Older adults also showed a shift from frontal and parietal mediators towards parietal and occipital mediators, raising the question of whether the gateway through which genetic variants modulate intelligence changes during the lifespan, to functionally compensate for aging processes ^16,65^. While the lack of frontal mediators in older adults may seem surprising at first glance given the well-established posterior-anterior shift in aging model (PASA)^66^, it is important to note that PASA has been observed in task-based studies. Interestingly, a recent study successfully replicated PASA via resting-state EEG ^67^ . In line with our results, the authors additionally found reduced intra-area frontal connectivity and an increased inter-area connectivity in older adults compared to young adults. This reduction in segregation fits our assumption that with age, slow oscillations may promote broader, cross-area communication, reflecting neural dedifferentiation. Notably, V1, cuneus, and the SPL remained consistent mediators across the lifespan. These regions are all involved in sensory processing or integration suggesting that sensory functions are crucial for maintaining cognitive ability. The pathway through which genetics shape behavior might be as dynamic as the brain itself, always striving for the most functional connection and our findings suggest that functional connectivity properties of sensory rather than frontal regions might underpin lifelong intelligenc*e*.

One limitation is that our dataset does not allow to investigate the association between SNPs, functional connectivity, and intelligence across the lifespan. Future studies should investigate potential reorganization of brain networks over time by means of longitudinal designs. It remains unclear whether intelligence in older adults relies on the same networks as in younger subjects. Research on the P-FIT model did not differentiate between age groups, and prior studies typically involved younger participants ^14,72–75^. Future research could benefit from a multi-modal approach, as Thiele et al. ^76^ have shown that overlapping as well as separate information is captured by different metrics. The complexity of intrinsic brain dynamics might be not fully captured by a narrow set of graph metrics, and a broader range of parameters, including complexity values, microstate characteristics ^76^ or other graph metrics such as modularity ^77^, might shed further light on the complex yet fascinating association between the genome, the brain, and human intelligence.

To the best of our knowledge this study is the first to investigate the mediating effects of EEG-derived graph metrics in specific brain regions on the association between genetic variation and intelligence in two different age groups. Our findings suggest that functional theta connectivity of frontal, parietal, temporal, and occipital areas provides a missing piece in the link between genetic variation and general intelligence. Different mediating effects between young and older adults raise the question of whether genetic pathways for intelligence evolve over the lifespan, highlighting the need for future research. Our findings are a crucial step forward in decoding the neurogenetic underpinnings of intelligence, as they identify specific electrophysiological networks that relate polygenic variation to intelligence.

## Methods

### Participants

The data stem from the ongoing large-sample cohort study *Dortmund Vital Study* (Clinicaltrials.gov: NCT05155397) conducted by the Leibniz Research Centre for Working Environment and Human Factors at the Technical University Dortmund (IfADo) that investigates the age-dependent development of cognitive functions in adult humans ^78^. The sample includes healthy subjects from any educational level, coming from a Western society. The health criteria were defined in such a way that smoking, alcohol consumption, being overweight, and/or a history of disease without severe symptoms did not exclude subjects from participating in the study. This approach of using more liberal participation criteria increases the representativeness of the sample^78^. Participants were recruited by newspaper ads, public media, companies, and public institutions. In total 599 participants were available. After excluding participants who lacked a major part of data subtests for the calculation of *g* (see 2.3. Computation of *g* factor), calculation of PGS_GI_ (see 2.4. Genotyping and polygenic scores), excluding participants without EEG data and controlling for outliers (see 2.7. Statistical analysis) the final data set included 434 participants (mean age: 44.4 years, SD: 14 years, age range: 20 – 70 years, 275 women). Before any data acquisition, all participants gave written informed consent, and the study conformed to the Declaration of Helsinki. In addition, it was approved by the local Ethic Committee of IfADo (A93-1).

To test whether associations between PGS, rsEEG and *g* vary between age groups, we analyzed the data for young adults and older adults separately. The younger sample comprised 199 participants (mean age: 31.23 years, SD: 6.73 years, age range: 20 – 40 years, 131 women) and the older sample comprised 235 participants (mean age: 55.65 years, SD: 6.84 years, age range: 40 – 70 years, 144 women). The cut-off value of 40 years was chosen based on the age distribution of the sample (see Supplementary Figure S1). The subgroups were analyzed in the same manner. Results for the full sample are shown in Supplementary Figures S3-S5.

### Cognitive measures

Participants completed an extensive cognitive test battery covering the essential aspects of intelligence. The test battery is described in the following and was used to calculate *g*, the factor of general intelligence (See 2.3. Computation of *g* factor). For detailed description of the tests please see Gajewski et al. ^79^.

### The Verbal Learning and Memory Task

The Verbal Learning and Memory Task (VLMT) ^80^ tests verbal declarative episodic memory. Here the experimenter presents 15 words from a learning list by reading them out loud to the participant. This is done five times and after every presentation, the participants have to reproduce the words from the learning list. Then, an interference word list with 15 new words is presented and has to be reproduced. Right after this, the original learning list has to be reproduced once more. 30 minutes later the participants are asked to repeat the learning list again. Lastly, participants are presented with a recall list, containing the items from the learning list, the interference list, and 20 new words. The items are presented orally, and the participants have to recognize the words belonging to the learning list. Scores are the total number of words from the learning list that have been reproduced in the five trials (VLMT_1_5) and the number of recognized words from the recognition list minus errors (VLMT_R-E).

#### D2-R

The D2-R measures attentional endurance and processing speed ^81^. Participants are presented with 14 lines consisting of 47 characters each. The characters are the letters “d” and “p” with one to four dashes above and/or below each letter. Participants have 20 seconds per line to cross out the target stimulus, which is a “d” with two dashes. The test score is the total number of correctly crossed-out target stimuli.

### Stroop Test

The color-word interference test ^82^ measures the processing speed and inhibition of incongruent information ^83^. Firstly, color words (e.g. “blue”) are presented in black and have to be read as quickly as possible. Secondly, colored bars are presented, and the color has to be named as quickly as possible (Stroop_2). Thirdly, participants see color words that are printed in a color that does not match the color word (e.g. the word “blue” printed in green). The participants have to indicate the color the word is written in (Stroop_3). Each of the three blocks consists of 36 items.

### Trail Making Test

The Trail Making Test (TMT) consists of two different tests ^84^. In TMT-A, participants have to connect the numbers 1 to 25 in ascending order. In TMT-B, the numbers 1 to 13 and the letters A to L have to be connected alternately in ascending order. While TMT-A measures processing speed, TMT-B measures parallel processing and the ability to switch between tasks.

### Digit Span

The Digit Span test measures memory span and working memory ^83^. First, the experimenter presents a series of digits with increasing length orally. The participants have to repeat the series of digits in the correct order (DS_f). Reproduction of two series of the same length counts as a correct response, the maximum length of the series is nine digits.

Second, the presented series of digits must be reproduced in reversed order. The maximum length of series is eight digits (DS_b). Again, the reproduction of two series of the same length counts as a correct response. The sum of DS_f and DS_b comprises the score DS_total.

### Multiple Choice Vocabulary Test

The Multiple Choice Vocabulary Test (MWT-B) measures verbal knowledge as a component of crystalized intelligence ^85^. Participants are exposed to 37 items with five words each. Only one of the five words is a meaningful word. The participants have to mark the meaningful word. The items get increasingly difficult over time. The total count of correctly marked words comprises the score.

### Word Fluency

Word fluency (WF) is subtest six from the performance testing system (Leistungsprüfsystem (LPS)) that measures verbal processing speed and cognitive flexibility ^86^. The participants are given three letters (F, K, R) and are asked to name as many words starting with these letters as possible. The participants have one minute for every letter and must write the words down (WF_W). In a second test, the participants say the words with the initial letters (B, F, L) (WF_S). The spoken version of the WF test was introduced to reduce age-related differences in writing speed. The test scores are the total number of non-repeated words written or spoken.

### Logical reasoning

The third subtest of the performance test system (LPS-3) measures logical reasoning, i.e., the ability to think logically as a component of fluid intelligence. The aim is to indicate the incongruent element in each row of eight logically arranged symbols. The highest score to be achieved is 40. The respondent was given five minutes to complete the test. The number of correctly indicated rows was used as dependent variable.

### Mental rotation

The seventh subtest of the performance test system (LPS-7) requires spatial rotation of letters in the plane, i.e., an ability attributed to fluid intelligence. The task consists of crossing out those letters that are recognized as mirror images. The time limit for this subtest is two minutes. A maximum of 40 recognized symbols can be achieved. The number of correctly crossed out letters was used as dependent variable for performance evaluation.

### Computation of g factor

The described cognitive tests have been used to generate the *g*-factor. After excluding all participants with missing values in one or more of the included variables (VLMT_1_5, VLMT_R-E, D2-R, Stroop_2, Stroop_3, TMT-A, TMT-B, DS_total, LPS_3, LPS_7, DS_f, DS_b, MWT, WF_S, WF_W), 520 participants remained for the factor analysis. To extract residuals from the test scores, individual regression analyses were calculated with age, sex, age*sex, age^2^, and age^2^*sex. Age^2^ was added to control for quadratic relations with age ^87^. Subsequently, the residuals were z-standardized (M = 0, SD = 1).

First, an exploratory factor analysis (EFA) with the estimation method “minimum residual” and “oblimin” factor rotation (an oblique, non-orthogonal rotation method) was used to assess the *g*-factor. The EFA yielded four factors, interpreted as verbal memory (including VLMT_1_5, VLMT_R-E), attention (including D2-R, Stroop_2, Stroop_3, TMT-A, TMT-B, DS_total, LPS_3, LPS_7), working memory (including DS_f, DS_b, MWT), and verbal fluency (including MWT, WF_S, WF_W).

Second, we employed a second-order confirmatory factor analysis. Supplementary Figure S2 depicts the postulated confirmatory factor model of *g*, including z-standardized factor loadings and covariances between subtests. This model was used to calculate *g* for every participant. Model fit was determined via multiple indices. The chi-squared statistic, assessing if the difference between the variance-covariance matrix implied by the model and the observed variance-covariance model is zero ^88^, reached statistical significance (X^2^(82) = 194.529, p < .001). However, this alone does not indicate a poor fit, as X^2^ is heavily influenced by sample size and can reach significance in large sample sizes even though the model fit is good ^89,90^. Root Mean Square Error of Approximation (RMSEA) and Standardized Root Mean Square Residual (SRMR) both indicate a good model fit (RMSEA = 0.051, SRMR = 0.042), while the Comparative Fit Index (CFI) and the Tucker-Lewis Index (TLI) closely miss the threshold of 0.97 (CFI = 0.950, TLI = 0.936).

### Genotyping and polygenic scores

Blood samples were collected from the participants in EDTA blood test tubes. DNA isolation was carried out on 5 ml of whole blood using the QIAamp DNA Blood Maxi Kit (Qiagen, Hilden, Germany). Quality of the DNA samples was ensured via NanoDrop (260/280 nm ratio) (Thermo Fisher Scientific Inc), and samples were stored at -20 °C. Illumina’s Infinium Global Screening Array 3.0. with MDD and Psych content (Illumina, San Diego, CA, USA) were used for genotyping. Genotyped SNPs with a minor allele frequency (MAF) lower than 0.05, deviating from Hardy-Weinberg equilibrium (HWE) with *p*<1*10^-6^ and missingness higher than 2%, were excluded. Participants showing sex mismatch, an SNP-missingness rate higher than 2%, and a heterozygosity rate higher than |0.2| were excluded. Furthermore, genetic relatedness filtering was carried out on an SNP-subset showing high genotyping quality (HWE *p*>0.02, MAF>0.2, SNP-missingness of 0%) and pruned for linkage disequilibrium (r^2^=0.1). Pairs of cryptically related subjects with a pi hat value greater than 0.2 were applied to exclude subjects at random. Principal components (PCs) were computed to control for population stratification. Any participant deviating by more than |4.5| standard deviations from the mean on at least one of the first 20 PCs was also excluded. All filtering steps were performed using PLINK 1.9 ^91^.

The samples’ filtered genotype data was submitted for imputation to the Michigan Imputation Server ^92^ using the phase 3v5 (hg19) European population of the 1000 Genomes Project as a reference panel, and an R^2^ filter of 0.3. We chose Eagle 2.4 for phasing and Minimac4 for imputation. After a final MAF<0.05 filtering step, 6,014,815 SNPs remained available for analysis.

We calculated genome-wide polygenic scores (PGS) for all participants, using publicly available summary statistics of general intelligence ^11^. General intelligence PGS (PGS_GI_) was calculated by adding together the weighted sum of all trait-associated alleles across SNPs with a high imputation quality in the summary statistics (INFO score > 0.9) present in both the summary statistics and in the imputed sample using PRSice2.3.3 ^12^. Specifically, we computed best-fit PGS_GI_ using standard settings by carrying out multiple linear regression iteratively at different SNP-*p*-value thresholds ranging from 5*10^-5^ to 1 (increasing the threshold each time by 5*10^-5^) to predict *g*. While the PGS_GI_ served as predictors, age, sex, as well as the first four genetic PCs were entered as covariates. The PGS_GI_ with the greatest predictive power (i.e., with the largest R^2^ increment when entered a model containing only the covariates) were chosen for further analysis. So, the resulting PGS explained the most phenotypical variance among all tested models. Since this approach involves parameter optimization, the best-fit model’s *p*-value is overfitted. To account for this, we used PRSice-2’s permutation-based approach to correct for multiple tests that consist of repeating the above procedure 10,000 times but shuffling the phenotype data with each iteration. PGS_GI_ best predicted *g*’s variance (4%, *p*_corrected_ < 0.001) in our sample at a p-value threshold of 0.328.

### EEG recordings and pre-processing

The Dortmund Vital Study comprises a battery of mental tasks that are distributed over two different days, with each testing session lasting around two hours, and involving EEG recordings ^78^. Before and after each test session resting-state EEG is recorded. Each recording consists of two minutes resting-state EEG with eyes closed and with eyes open each, and EEG recording session 1 and 2 are always around two hours apart. For the present analysis, only the two EEG data sets recorded on the first testing day were used, as the EEG recordings on the second day used different EEG recording systems ^93^. To keep the conditions consistent with existing literature on resting-state EEG functional connectivity and human intelligence, only the eyes-closed condition was used for further analyses ^28–30^. A total of 438 of the participants with genetic and behavioral data also had complete resting-state EEG data. The EEG data employed in this study were recorded using a 64-channel actiCap system and a BrainAmp DC-amplifier (Brain Products GmbH; Munich, Germany) with a 1000 Hz sampling rate. An online 250 Hz low-pass filter was applied and FCz was used as an online reference. Electrodes were placed in line with the 10-20 system and impedances were < 10 kΩ^78^. EEG data preprocessing was performed using custom MATLAB scripts with EEGLab ^94^. The recordings of all resting-state conditions and sessions were merged and subsequently resampled at 200 Hz. The data was filtered using a Butterworth bandpass filter ranging from 1 to 30 Hz with the order 4. Channels with insufficient data quality were detected and rejected based on kurtosis and probability criteria. After that, excluded channels were interpolated and all data were re-referenced to the common average reference. The continuous data were then segmented into epochs of eight seconds in length with an overlap of 50 %. Epochs with bad data quality were identified and excluded from further analysis using EEGLab’s automated rejection function with default parameters. An individual component analysis (ICA) was performed, and independent components (ICs) reflecting artifacts were identified using ICLabel ^95^ and subsequently removed. ICs were regarded as reflecting artifacts when ICLabel classified an IC with a probability of less than 0.3 to reflect brain activity, as well as when an IC was classified with a probability of more than 0.3 to reflect ocular activity. The inter-areal functional connectivity was calculated based on the brain activity at reconstructed sources using MNE python ^96^. Since no individual T1 MRI scans and electrode positions were available, MNE templates were used to set up the source space, the boundary element model, and the electrode montage. Based on these, the forward model was then computed. The inverse operator was calculated for each dataset using an individual noise covariance matrix. Source reconstruction was performed using the dSPM method ^97^, a minimum norm approach that has previously been used for the source reconstruction of resting-state data ^29,30^. The PALS-B12 atlas^98^ was used for the parcellation of the sources in 41 regions of interest (ROI) per hemisphere that correspond to Brodmann areas (Brodmann areas not included in the PALS-B12 atlas: BA12, BA13, BA14, BA15, BA16, BA34, BA48, BA49, BA50, BA51, BA52). Source activation for these ROIs was extracted using MNEs “extract_label_time_cource” method with the “mean_flip” mode. All-to-all inter-ROI functional connectivity was calculated as the spectral coherence for the delta (2 to 3 Hz), theta (4 to 7 Hz), low alpha (8 to 10 Hz), high alpha (11 to 13 Hz), and beta (16 to 30 Hz) frequency range using MNEs “spectral_connectivity_time” method. Alpha was divided into low alpha and high alpha given that past studies repeatedly revealed correlations between alpha and intelligence, especially within the upper alpha band ^28,99–101^. Whereas splitting alpha into sub-bands has proven to deliver additional information in terms of cognitive abilities, many studies have not demonstrated significant associations between intelligence and sub-bands of other frequencies, such as beta or gamma, during resting-state EEG ^28–30^. Hence, we have chosen to only divide alpha. For each ROI-to-ROI spectral coherence, the mean spectral coherence of 30 segments was estimated. 30 segments of on average 8 seconds and a sampling rate of 200 Hz result in a total of approximately 48000 data points, which is notably higher if compared to an fMRI dataset of the same length.

### Graph metrics

Global and nodal efficiency was calculated using the Brain Connectivity Toolbox and in-house MATLAB code ^102^. We constructed two 82 x 82 resting-state networks per frequency band (delta, theta, low alpha, high alpha, beta), one for the first eyes-closed recording and one for the second eyes-closed recording (i.e., before and after the participants performed the two-hour battery of mental tasks). As the conditions during and between the two resting-state recordings were the same for all subjects, we combined these two matrices into one 82 x 82 matrix by calculating the mean coherence of each connection. Thus, we analyzed five 82 x 82 networks. To prune redundant and weak connections we employed Holm-Bonferroni pruning with a threshold of 0 for each network (□ = 0.01, one-tailed) as described by Ivković et al. ^103^. This pruning procedure prevents some of the drawbacks of fixed thresholds like including spurious connections or excluding important connections. Here we used the variance of all weights in the upper triangle of the matrix of all participants to test if an edge weight is a spurious connection or not. This was done for all connections. For example, a vector containing participants’ edge weights for the edge between left BA10 and left BA11 was tested against zero and removed from the network if it did not differ significantly from zero. This was done for all edges and led to no connections being removed from the networks.

We used MATLAB R2021b and the Brain Connectivity Toolbox to compute global efficiency (*E*), global clustering (*C*), nodal efficiency (*E_i_*), and local clustering (*C_i_*) indices. Global efficiency *E* quantifies how efficiently – fast but also using little energy – the brain areas communicate throughout the brain ^15^. Large edge weight and short path length lead to an increase in global and nodal efficiency. A shortest path is defined by the minimal number of edges that are needed to connect two nodes within a network. The distance matrix *d* contains all shortest paths between all node pairs. This matrix is created by calculating the inverse of the weighted adjacency matrix and running Dijkstra’s algorithm ^104^. The efficiency of a single brain area is called the nodal efficiency *E_i_*. It reflects the average inverse shortest path length between a node *i* and all other nodes *j* in the network *G.* Global efficiency reflects the average inverse shortest path length between all nodes *i* and all other nodes *j* in *G*:

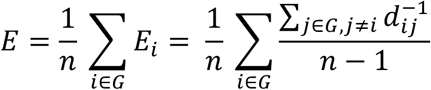

The global clustering coefficient *C* is a measure to quantify the “cliquishness” of a network and reflects the network’s local connection segregation ^102^. The local clustering coefficient *C_i_* reflects the probability that two randomly selected neighbors of node *i* are also neighbors of each other. *C_i_* is calculated by dividing the real connections between a node’s neighbors by all possible connections. The global clustering coefficient *C* is defined as the mean of all local clustering coefficients.

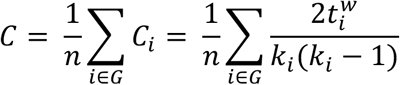

### Statistics and Reproducibility

All statistical analyses were performed using RStudio 1.3.1093 ^105^ and R version 4.1.0 ^106^. Participants that deviated more than three interquartile ranges from the sample’s global efficiency, global clustering coefficient, or *g* were classified as outliers and removed from the analyses. Four participants were excluded by this step resulting in a final sample size of 434 participants.

### Partial correlations

We calculated partial correlations between *g* and the PGS_GI_ and the brain metrics global efficiency and global clustering coefficient (two-sided). For this, we used the *partial.cor* function from *RcmdrMisc* ^107^. Control variables were age and sex.

### Global mediation analyses

Mediation models were calculated using the *lavaan* package ^108^. The dependent variable was *g*, and the independent variable was the PGS_GI_. Respective mediators were global efficiency or global clustering. We controlled for age, sex, and the first four principal components of the population stratification.

### Brain area-specific mediation via elastic-net regression

The next step was to investigate if there is a subset of brain areas whose connectivity may act as a mediator of the association between PGS_GI_ and *g*. For this, we applied exploratory mediation analysis by regularization (*xmed*) ^109,110^. Since this method employed regularization instead of frequentist methods, there are no p-values to determine whether or not a brain area can be considered a viable mediator ^111^. Regularization means that a penalty term is put onto effect sizes which shrinks small effect sizes to zero. All effect sizes that remain non-zero after regularization are viable mediators. To prevent overfitting, the penalty is calculated using *k*-fold cross-validation. Here, the sample is split into k subsamples. One subsample is used as the testing set while all other subsamples serve as training data. This is repeated *k* times, with all subsamples serving as testing sets once. As this method is calculating a mediation, two different elastic-net models are calculated, one for path *a* (brain ∼ PGS_GI_) and one for path *b* (g ∼ brain). The mediation coefficient is obtained by multiplying *a* and *b*. Thus, both the association between brain and PGS_GI_ as well as brain and *g* must be classified as non-zero for the mediation coefficient to be non-zero. All non-zero mediation coefficients are selected as mediators. As the penalty pushed all effect sizes close to zero, the effect sizes derived by elastic-net are biased. Thus, we followed the suggestion by Serang & Jacobucci ^110^ and re-estimated the effect sizes of selected mediators using *lavaan*. We used elastic-net regression as it is a combination of lasso and ridge regression and is suitable if one does not have a clear expectation regarding all mediating variables. Regularized elastic-net regression has already been successfully applied in recent neuroimaging-based intelligence research ^26,40,112^.

### Specific mediation models

We used the *xmed* function from the *regsem* package 1.9.3. ^113^ for all brain area-specific mediation analyses. For the analysis in this manuscript, the cross-validation fold was set to k = *10* (default). Control variables were sex, age, and the first four principal components of the population stratification. The threshold for detecting non-zero mediation effects was set to 0.001 and □ was set to 0.5, corresponding to a full elastic-net penalty. To investigate the effect of paths *a* and *b* separately, we defined a threshold of 0.01 for identifying non-zero effects. The threshold for paths *a* and *b* were set higher as the product of the two is smaller since both are always below one. The dependent variable was *g*, and the independent variable was the PGS_GI_. Respective mediators were nodal efficiency or local clustering coefficient of 41 cortical areas in the left and 41 cortical areas in the right hemisphere. We calculated one mediation model per graph metric and per frequency band, resulting in ten mediation models in total: PGS_GI_ – delta nodal efficiency – *g*, PGS_GI_ – theta nodal efficiency – *g*, PGS_GI_ – low alpha nodal efficiency – *g*, PGS_GI_ – high alpha nodal efficiency – *g*, PGS_GI_ – beta nodal efficiency – *g*, PGS_GI_ – delta local clustering – *g*, PGS_GI_ – theta local clustering – *g*, PGS_GI_ – low alpha local clustering – *g*, PGS_GI_ – high alpha local clustering – *g*, PGS_GI_ – beta local clustering – *g*.

### Test-retest reliability of graph metrics

As we had two EEG recordings taken approximately 2 hours apart from each other, we used both the first eyes-closed recordings (EC1) and the second eyes-closed recording (EC2) to investigate test-retest reliability. We calculated the ICC (3,1) with a two-way mixed effect model ^93^. First, we calculated the ICC of the global efficiency and global clustering coefficient of the five frequency bands. Second, we calculated the ICC of nodal efficiency and local clustering of the five frequency bands for all 82 cortical areas.

## Supporting information

Supplemental Figures and Supplemental Results

## Acknowledgments

The authors are grateful to Tobias Blanke for technical support and to the lab staff, in particular to Claudia Brockhaus, Pia Deltenre, Barbara Foschi, Carola Reiffen, Silke Joiko, Christiane Westedt and their team of student assistants for their help with data acquisition. The study is endorsed by the German Center for Mental Health (DZPG). The Dortmund Vital Study is funded by the institute’s budget. Thus, the study design, collection, management, analysis, interpretation of data, writing of the report, and the decision to submit the report for publication is not influenced or biased by any sponsor.

## Author Contributions

E.G., E.W. and R.K. conceived the project. E.G., R.E., D.M., S.G., P.D.G., E.W., M.B., J.G.H., C.W., and M.A.N. designed the project and supervised the experiments. S.G., P.D.G. and C.F. performed data collection. J.D., D.M., C.S. and E.G. planned and performed statistical data analyses. J.R., J.S.P., F.S., S.O. and R.K. planned and performed genetic data analysis and supported manuscript drafting. S.A. and D.S. planned and performed EEG data analyses and supported manuscript drafting. D.M., R.E., C.S. and E.G. wrote the manuscript, with substantial support from all authors. All authors discussed the results and have contributed significantly to the final submitted version of the manuscript.

## Data Availability Statement

The data and R code that support the findings of this study are available from the corresponding author upon reasonable request or can be downloaded from an Open Science Framework repository [https://osf.io/c9bwz/].

## Code Availability Statement

The MATLAB and Python code used for pre-processing, source reconstruction, and the calculation of the connectivity measures is available at the GitHub repository [https://github.com/stefanarnau/EEG_resting_state_network_intelligence].

