## Supplemental Figures and Supplemental Results for "Electrophysiological resting-state signatures link polygenic scores to general intelligence"

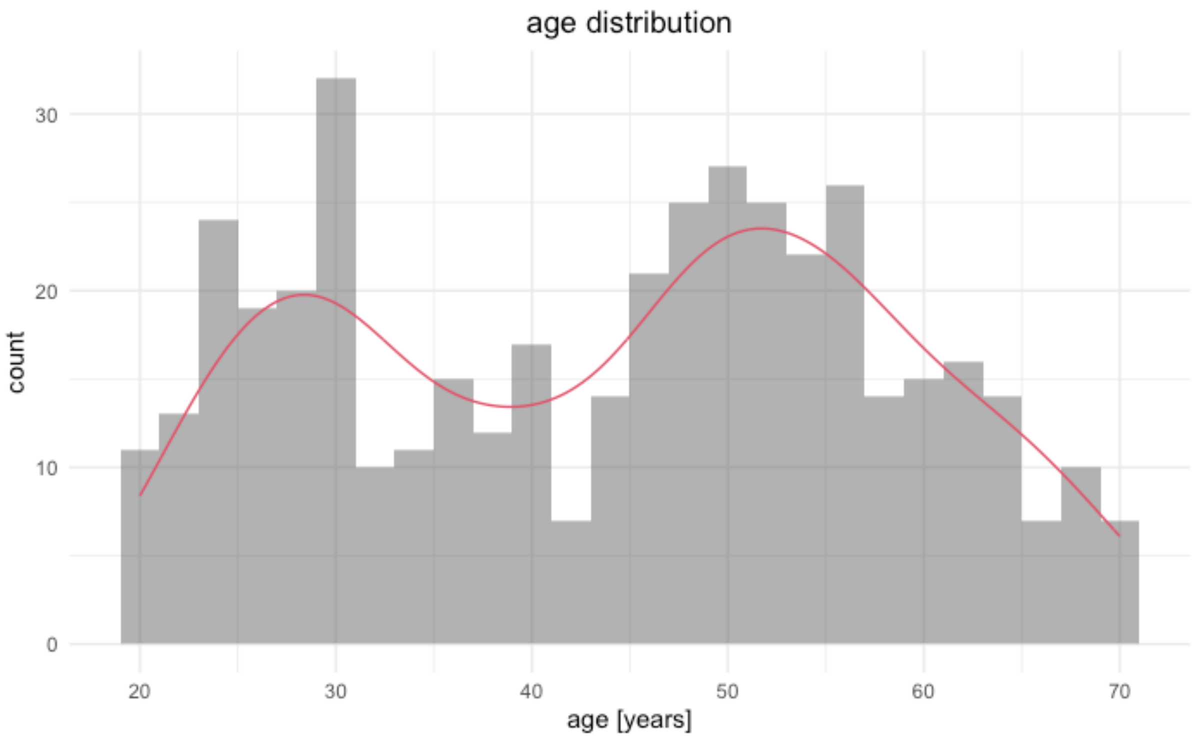


**Supplementary Figure S1.** Age distribution in the whole sample.

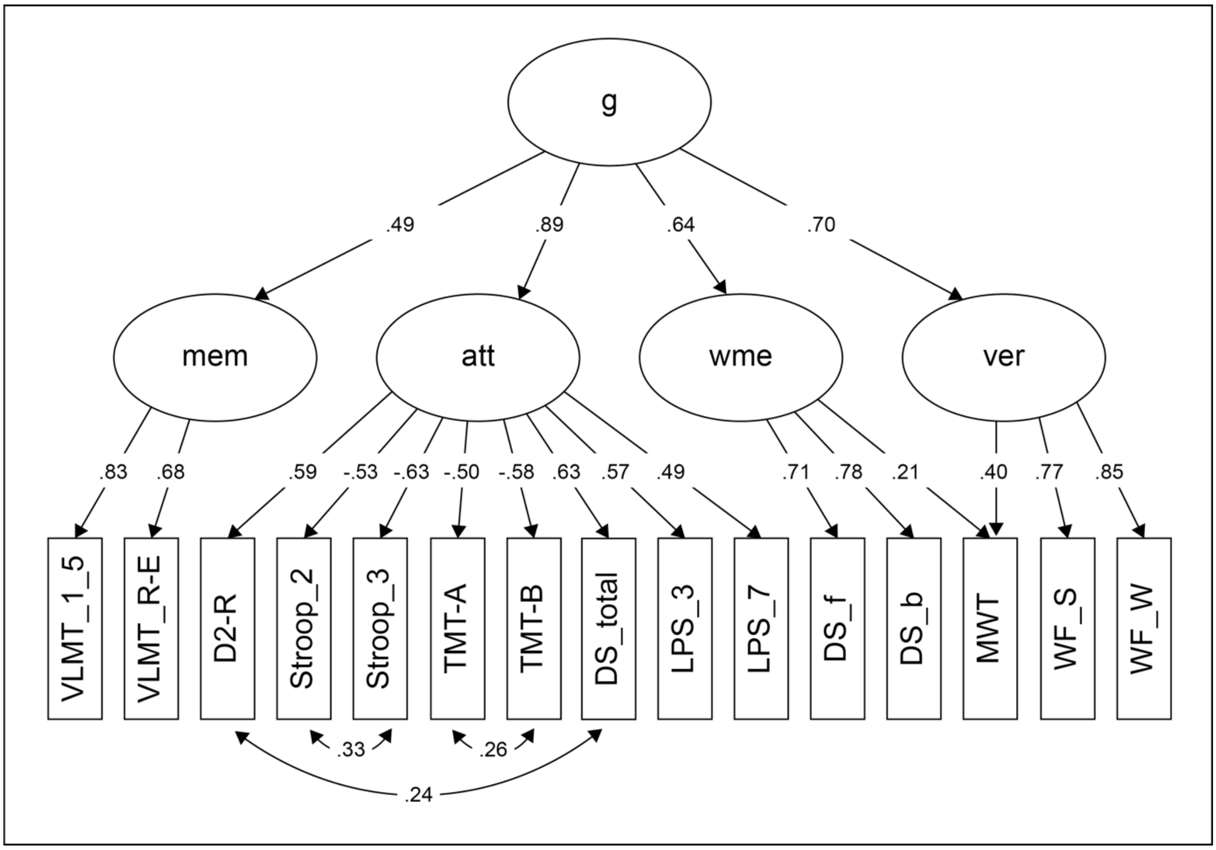


**Supplementary Figure S2**. Confirmatory factor analytic model. *g* = general factor of intelligence, mem = verbal memory, att = attention, wme = working memory, ver = verbal fluency. VLMT = Verbal Learning and Memory Task, TMT = Trail Making Test, DS = Digit Span, LPS = Leistungsprüfsystem, MWT = Multiple Choice Vocabulary Test, WF = Word Fluency.

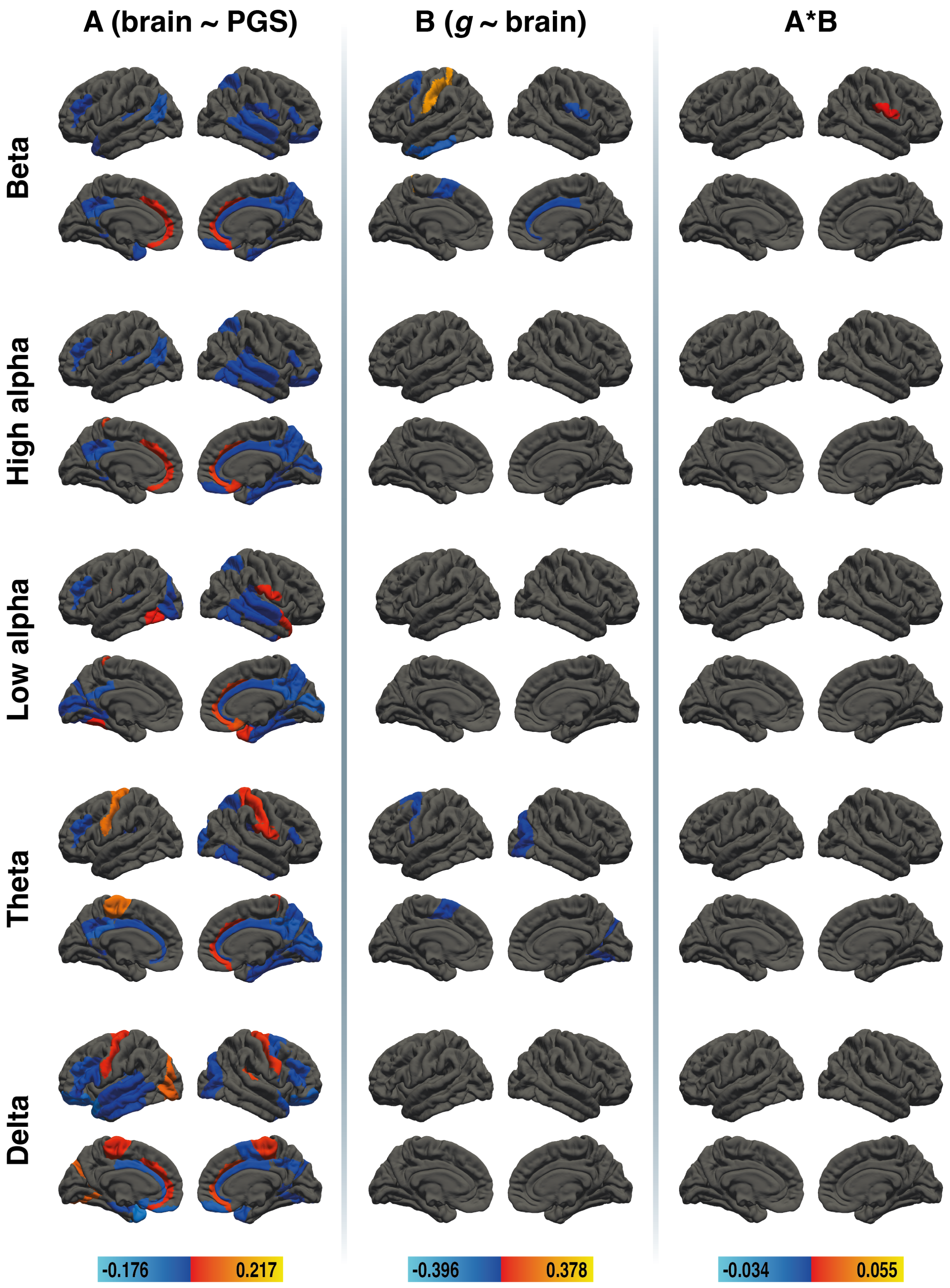


**Supplementary Figure S3.** Results of the region-specific mediation analysis via elastic-net regression (nodal efficiency). Respective mediators were 41 cortical areas in the left and 41 cortical areas in the right hemisphere of five different frequency bands (delta, theta, low alpha, high alpha, and beta, from the bottom to the top). The figure shows the results from path a analysis (brain ~ PGS), path b analysis (g ~ brain), and the mediation effect (from left to right). Brain surfaces are shown in lateral and sagittal view, for the left and right hemisphere. Positive effects are depicted in red and yellow, negative effects are depicted in blue.

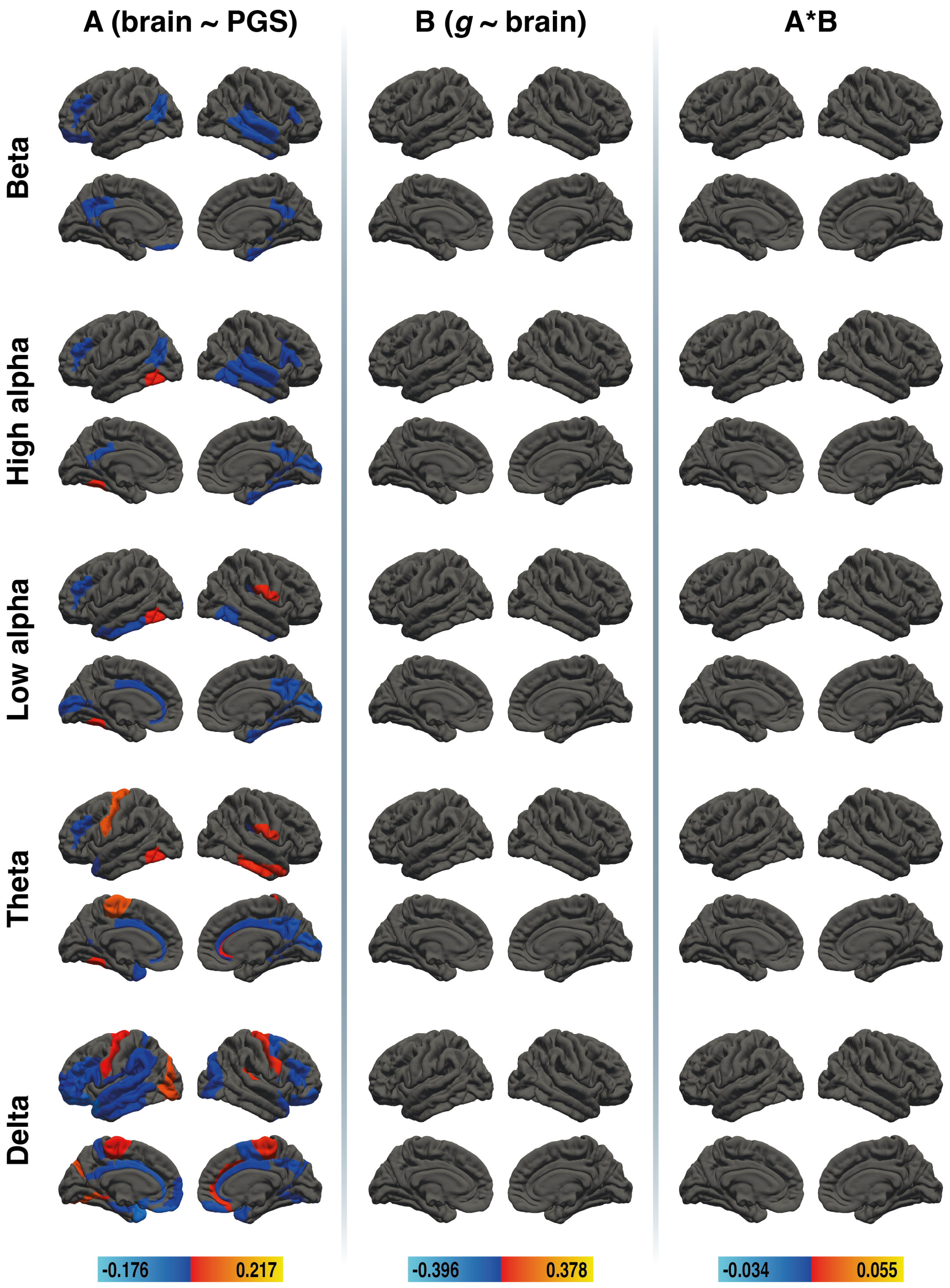

**Supplementary Figure S4.** Results of the region-specific mediation analysis via elastic-net regression (local clustering). Respective mediators were 41 cortical areas in the left and 41 cortical areas in the right hemisphere of five different frequency bands (delta, theta, low alpha, high alpha, and beta, from the bottom to the top). The figure shows the results from path a analysis (brain ~ PGS), path b analysis (g ~ brain), and the mediation effect (from left to right). Brain surfaces are shown in lateral and sagittal view, for the left and right hemisphere. Positive effects are depicted in red and yellow, negative effects are depicted in blue.

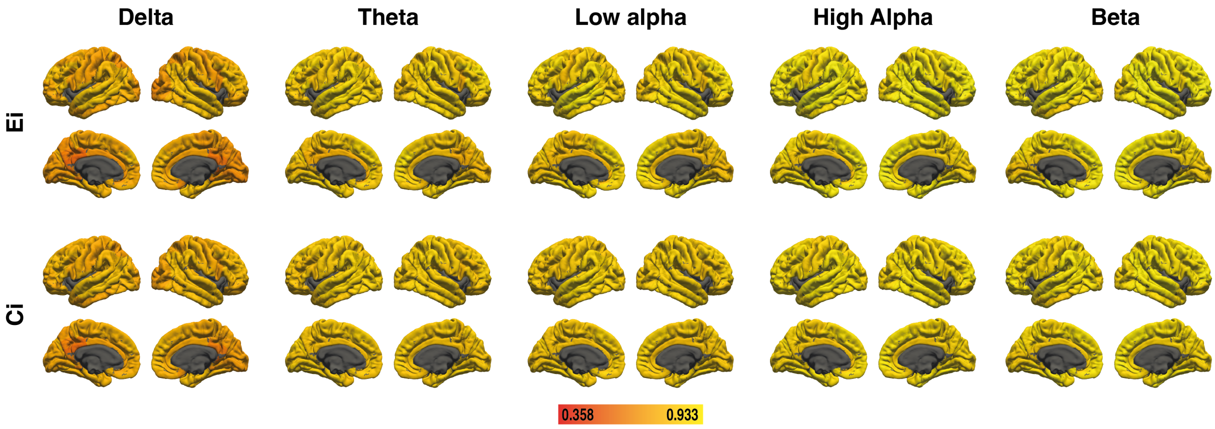


**Supplementary Figure S5.** Test-retest reliability of nodal efficiency ($E_i$) and local clustering ($C_i$) as measured by ICC. Lighter colors indicate higher reliability.

### Supplementary Results

#### Results for the complete sample

##### *Partial correlations*

Partial correlations between *g*, global efficiency, and global clustering coefficient are shown in Supplementary Table S1. Partial correlations between $PGS_{GI}$, global efficiency, and global clustering coefficient are shown in Supplementary Table S2. No correlation reached statistical significance.

##### *Global mediation analysis*

Results of the global mediation analysis are shown in Supplementary Table S3. None of the global mediation analysis reached statistical significance. Even though partial correlations conducted prior to this analysis did not reveal a significant effect, the global mediation analysis were still conducted, as statistical significance of paths a and b alone are not a requirement for a significant mediation effect (Hayes 2018; Zhao et al. 2010).

##### *Brain-area specific mediation analyses*

*Nodal efficiency*

Results for the brain-area specific mediation analysis for the full sample for nodal efficiency are depicted in Supplementary Figure S3.

*Beta*

$PGS_{GI}$ were associated with beta-frequency nodal efficiency in 25 cortical areas. 10 areas were located on the left hemisphere, 2 showed a positive association, 8 showed a negative association. 15 areas were located on the right hemisphere, 1 showed a positive association, 14 showed a negative association. There were six areas whose beta-frequency nodal efficiency was associated with *g*, namely left BA2, BA21 and BA6 as well as right BA24, BA30 and BA43. Left BA2 and right BA30 showed positive associations with *g*, while left BA21 and BA6 and right BA24 and BA43 showed negative associations. The beta-frequency nodal efficiency of right BA30 and BA43 were identified as mediators of the effect of $PGS_{GI}$ on *g*. Right BA30 showed a negative mediation effect, which is due to a negative association of BA30's beta-frequency nodal efficiency being negatively associated with $PGS_{GI}$ and positively associated with *g*. Right BA43 showed a positive mediation effect, because BA43's beta-frequency nodal efficiency was negatively associated with both *g* and $PGS_{GI}$.

*High alpha*

$PGS_{GI}$ were associated with high alpha-frequency nodal efficiency in 30 cortical areas. 10 areas were located on the left hemisphere, 3 showed a positive association, 7 showed a negative association. 20 areas were located on the right hemisphere, 2 showed a positive association, 18 showed a negative association. There were no areas whose high alpha-frequency nodal efficiency was associated with *g* and thus no selected mediators.

*Low alpha*

$PGS_{GI}$ were associated with low alpha-frequency nodal efficiency in 32 cortical areas. 12 areas were located on the left hemisphere, 3 showed a positive association, 9 showed a negative association. 20 areas were located on the right hemisphere, 4 showed a positive association, 16 showed a negative association. There were no areas whose low alpha-frequency nodal efficiency was associated with *g* and thus no selected mediators.

*Theta*

$PGS_{GI}$ were associated with theta-frequency nodal efficiency in 29 cortical areas. 9 areas were located on the left hemisphere, 2 showed a positive association, 7 showed a negative association. 20 areas were located on the right hemisphere, 4 showed a positive association, 16 showed a negative association. There were two areas whose theta-frequency nodal efficiency was associated with *g*, namely right BA19 and left BA6. Both areas displayed a negative association with *g*, but showed no association with $PGS_{GI}$, thus they were no mediators.

*Delta*

$PGS_{GI}$ were associated with delta-frequency nodal efficiency in 30 cortical areas. 17 areas were located on the left hemisphere, 4 showed a positive association, 13 showed a negative association. 13 areas were located on the right hemisphere, 3 showed a positive association, 10 showed a negative association. There were no areas whose delta-frequency nodal efficiency was associated with *g* and thus no selected mediators.

*Local clustering*

Results for the brain-area specific mediation analysis for the full sample for local clustering are depicted in Supplementary Figure S4.

Beta

$PGS_{GI}$ were associated with low alpha-frequency local clustering in 16 cortical areas. 7 areas were located on the left hemisphere, 9 areas were located on the right hemisphere. All areas showed a negative association with $PGS_{GI}$. There were no areas whose beta-frequency local clustering was associated with *g* and thus no selected mediators.

High alpha

$PGS_{GI}$ were associated with high alpha-frequency local clustering in 19 cortical areas. 6 areas were located on the left hemisphere, 2 showed a positive association, 4 showed a negative association. 13 areas were located on the right hemisphere, all showed a negative association. There were no areas whose high alpha-frequency local clustering was associated with *g* and thus no selected mediators.

Low alpha

$PGS_{GI}$ were associated with low alpha-frequency local clustering in 18 cortical areas. 6 areas were located on the left hemisphere, 1 showed a positive association, 5 showed a negative association. 10 areas were located on the right hemisphere, 1 showed a positive association,

9 showed a negative association. There were no areas whose low alpha-frequency local clustering was associated with *g* and thus no selected mediators.

Theta

$PGS_{GI}$ were associated with theta-frequency local clustering in 19 cortical areas. 6 areas were located on the left hemisphere, 2 showed a positive association, 4 showed a negative association. 13 areas were located on the right hemisphere, 4 showed a positive association, 9 showed a negative association. There were no areas whose theta-frequency local clustering was associated with *g* and thus no selected mediators.

Delta

$PGS_{GI}$ were associated with delta-frequency local clustering in 33 cortical areas. 21 areas were located on the left hemisphere, 3 showed a positive association, 18 showed a negative association. 12 areas were located on the right hemisphere, 3 showed a positive association, 9 showed a negative association. There were no areas whose delta-frequency local clustering was associated with *g* and thus no selected mediators.

***Reliability***

Supplementary Table S17 shows the ICC of global efficiency and global clustering coefficient for all frequency bands for the full sample. The ICC of global efficiency of the delta, theta, and low alpha bands can be rated as good, while the ICC of global efficiency of the high alpha and beta band can be described as excellent [1] . Reliability of global clustering coefficient can be rated as good for all frequency bands.

Supplementary Figure S5 shows ICC of nodal efficiency and local clustering for 82 cortical areas. A complete list of areas and ICC effect sizes can be found in Supplementary Table S15 for young adults and in Supplementary Table S16 for older adults. The delta band was the only frequency where areas showed a poor reliability (ICC < 0.5), however, this was only the case for 8.5% of areas considering nodal efficiency and 2.4% of areas considering local clustering. The areas with poor reliability are primarily located around the posterior cingulate cortex and precuneus.

**Acknowledgments**

The authors are grateful to Tobias Blanke for technical support and to the lab staff, in particular to Claudia Brockhaus, Pia Deltenre, Barbara Foschi, Carola Reiffen, Silke Joiko, Christiane Westedt and their team of student assistants for their help with data acquisition. The study is endorsed by the German Center for Mental Health (DZPG).
